# Evaluation of Zoetis GeneMax Advantage genomic predictions in commercial *Bos taurus* Angus cattle

**DOI:** 10.1101/2020.10.23.353144

**Authors:** Brian. C. Arisman, Troy N. Rowan, Jordan M. Thomas, Harly J. Durbin, William R. Lamberson, David J. Patterson, Jared E. Decker

## Abstract

The GeneMax (GMX) Advantage test, developed by Zoetis, uses approximately 50,000 single nucleotide polymorphisms (SNP) to predict the genomic potential of a commercial Angus heifer. Genetic predictions are provided for Calving Ease Maternal, Weaning Weight, Heifer Pregnancy, Milk, Mature Weight, Dry Matter Intake, Carcass Weight, Marbling, Yield, and three economic selection indices. Test results can inform selection and culling decisions made by commercial beef cattle producers. To measure the accuracy of the trait predictions, data from commercial Angus females and their progeny at the University of Missouri Thompson Research Center were utilized to analyze weaning weight, milk, marbling, fat, ribeye area, and carcass weight. Progeny phenotypic data were matched to the respective dam, then the cow’s genomic predictions were compared to the calf’s age-adjusted phenotypes using correlation and linear model effect sizes. All tested GeneMax scores of the dam were significantly correlated with and predicted calf performance. Our predicted effect sizes, except for fat thickness, were similar to those reported by Zoetis. In conclusion, the GeneMax Advantage test accurately ranks animals based on their genetic merit and is an effective selection tool in commercial cowherds.

## INTRODUCTION

Prediction of quantitative traits using DNA markers in beef cattle was first commercialized in the 1990s. However, many of these tests relied on a small number of markers and failed validation trials (Van Eenennaam et al., 2007). Genomic prediction, the use of thousands of genome-wide DNA markers (Nejati-Javaremi et al., 1997; Meuwissen et al., 2001), has proven to be a much more efficacious strategy in driving genetic improvement (García-Ruiz et al., 2016; Taylor et al., 2016). Still, many farmers, ranchers (Weaber et al., 2014), extension professionals, and even academics question the effectiveness of genomic prediction in commercial beef cattle. Demonstrations of the ability of genomic tests to accurately predict genetic merit may encourage farmers and ranchers to adopt this technology and accelerate genetic progress in commercial herds. Our objective is to evaluate the effectiveness of the Zoetis GeneMax Advantage (genomic predictions designed for commercial heifers) in predicting the genetic merit of Angus cattle. We hypothesize that, because this test was built using principles of genomic prediction, the dam’s GeneMax scores will significantly predict her calves’ performance.

## MATERIALS AND METHODS

An Animal Care and Use Committee protocol is not necessary for this project as DNA samples were collected as part of routine animal production practices. However, the University of Missouri has a demonstration ACUC protocol, number 7491, which covers the procedures used in this research.

Cows were commercial Angus that were from a crossbred base that were straightbred Angus since 1995. Phenotypic and pedigree data were collected for commercial animals at the University of Missouri’s Thompson Research Center and entered into Angus Genetics Inc. Beef Improvement Records (BIR) program through AngusOnline.org. Pedigree, phenotype, and GeneMax Advantage score information were retrieved as Excel files from AngusOnline.org in June of 2018. Phenotypic records were collected from 1995 to 2018, however only calves born from 2003 to 2011 and from 2014 to 2018 had data reported to BIR and genotyped dams (**Figure 1**). Summary statistics are presented in **Table 1**. Excel files were read into R (Team, 2018) and similar files from different years were combined. Packages utilized included readr (Wickham et al., 2018b), ggplot2 (Wickham, 2016), tidyr (Wickham and Henry, 2018), dplyr (Wickham et al., 2018a), and stringr (Wickham, 2018).

**Figure 1.**
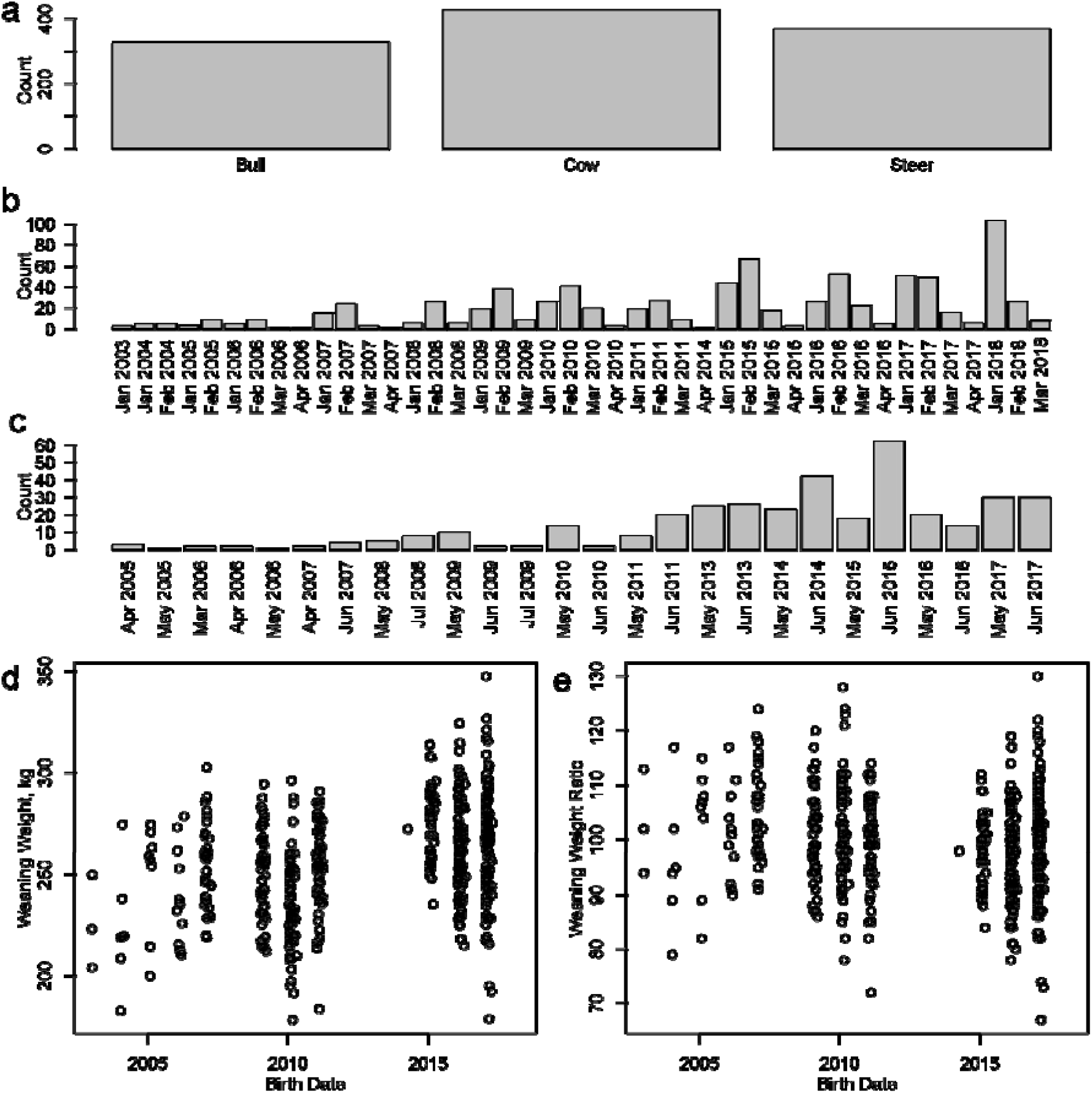
Graphical summary of data available for analysis. a) Counts of animals by sex. An animal can be counted as both a bull and a steer, for example if it was a bull at weaning but castrated prior to entering the feedlot. b) Counts of animals by birth month and year. c) Counts of animals by slaughter month and year. d) Weaning weight plotted against birth date. e) Weaning weight ratio plotted against birth date.

**Table 1.** Summary statistics of data used for GeneMax Advantage evaluation. For GeneMax scores, we report the number of animals with test results (Number) and the number of cows with calves with phenotypes (Number with matched calves).

|  | Number<br>(Number<br>with<br>matched<br>calves) | Mean | Standard<br>Deviation | Median | Minimum | Maximum | Skew | Kurtosis |
| --- | --- | --- | --- | --- | --- | --- | --- | --- |
| **Weaning**<br>**Weight,**<br>**kg** | 781 | 263.0 | 26.3 | 264.4 | 178.3 | 347.5 | -0.17 | 3.05 |
| **Weaning**<br>**Weight**<br>**Ratio** | 491 | 100.1 | 9.0 | 100 | 67 | 130 | -0.08 | 3.53 |
| **Marbling**<br>**Score** | 374 | 6.5 | 1.1 | 6.6 | 3.2 | 9.2 | -0.22 | 3.1 |
| **Hot**<br>**Carcass**<br>**Weight,**<br>**kg** | 376 | 398.7 | 40.3 | 401.9 | 249.5 | 504.9 | -0.62 | 3.93 |
| **Fat**<br>**Thickness,**<br>**cm** | 290 | 1.8 | 0.5 | 1.7 | 0.5 | 4.1 | 1.1 | 6.33 |
| **Ribeye**<br>**Area, cm<sup>2</sup>** | 374 | 84.8 | 9.3 | 84.5 | 57.4 | 112.9 | 0.12 | 2.87 |
| **WW**<br>**GMX**<br>**Score** | 554 (231) | 45.6 | 20.4 | 45.0 | 3 | 97 | 0.19 | 2.27 |
| **Milk**<br>**GMX**<br>**Score** | 554 (231) | 51.6 | 21.2 | 53.0 | 3 | 97 | -0.14 | 2.18 |
| **CW GMX**<br>**Score** | 554 (196) | 49.2 | 19.4 | 48 | 7 | 95 | 0.11 | 2.27 |
| **Marb**<br>**GMX**<br>**Score** | 555 (196) | 64.3 | 21.2 | 67 | 7 | 98 | -0.44 | 2.25 |
| **RE GMX**<br>**Score** | 556 (196) | 50.5 | 20.1 | 49 | 7 | 97 | 0.19 | 2.06 |
| **Fat GMX**<br>**Score** | 557 (188) | 43.3 | 19.9 | 42 | 3 | 93 | 0.26 | 2.18 |

For statistical analyses, the calf phenotype was compared with the dam’s GeneMax score (1 to 100 scale). For each of the traits, Pearson and Spearman correlations were calculated.

To adjust for additional factors, mixed models were used to evaluate the relationship between calf phenotype and dam’s GeneMax Score. We used the model:

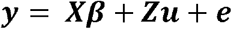

where, ***y*** is the phenotype of the calf; ***β*** are fixed effects of contemporary group, sex, and dam GMX Score; ***u*** is the random effect of sire, and ***e*** is the residual. Both ***u*** and ***e*** are ~*N*(0, *I*). For growth traits, the birth year of the calf was considered the contemporary group. For carcass traits, contemporary group was defined by the harvest date. There were 47 bulls who sired calves with weaning weight records, 37 bulls who sired calves with hot carcass weight, marbling, and ribeye area records, and 23 bulls who sired calves with fat thickness records. For each of the traits, four models were compared: a full model including dam GMX Score and the random effect of sire, a reduced model that did not include the effect of sire, a reduced model that did not include the effect of dam GMX Score, and a model with effect of sire and maternal grandsire (MGS). For each trait, a *χ*^2^ test was also run between the first and third models to determine the significance of the inclusion of the cow’s GeneMax Advantage score. A *χ*^2^ test was run between the third and fourth models to determine if the inclusion of the random effect of maternal grandsire was significant. Pseudo-*R*^2^ values were calculated using the ‘r.squaredLR’ function from the MuMIn package (Bartoń, 2022). Calf birth weights were not analyzed because GeneMax does not report a birth weight score or a calving ease direct score (only calving ease maternal). For weaning weight, phenotypes were pre-adjusted to 205-day weights and adjusted for age-of-dam effects by Angus Genetics Inc. Further, a model was also executed that included both Weaning Weight (WW) GMX Score and Milk GMX Score. Models containing 1) WW GMX Score, 2) Milk GMX Score, and 3) WW and Milk GMX Scores were compared to see which best fit the data. Because there is a trend over time for weaning weight phenotypes and we do not have a random sample of DNA tested cows, calves born in the early 2000s may have low weaning weights compared to calves born in later years (**Figure 1d**) but ranked high in their own contemporary group (weaning weight ratio, **Figure 1e**). In other words, entire contemporary groups were not analyzed, only calves of genotyped dams. Thus, some contemporary groups only had a small number of calves represented and the data have inherent selection bias. Therefore, we also measured the association between the dam’s GMX WW scores and GMX Milk scores with the calf’s weaning weight ratio (no contemporary group effect was included in these models).

Estimates of GeneMax Advantage score effects were retrieved from Zoetis technical bulletin GMX-00116 (Zoetis Genetics and Angus Genetics Inc., 2018). Effects were converted from Imperial to metric units, divided by 10 to represent a 1-point GeneMax score increase, and divided by 2 to change from molecular breeding values to expected progeny differences (transmitting abilities). Our GeneMax score effect estimates were compared to Zoetis’ published estimates using a two-tailed Z-test.

## Results

For each trait evaluated, the Pearson’s correlation and Spearman’s correlation between the calf’s phenotype and the dam’s corresponding GMX Score were statistically different from zero (**Table 2**). Further, from the six regression models, the dam’s GMX Score had a significant effect on the calf’s phenotype (**Table 3**). Except for fat thickness, our estimated effect sizes were not statistically different compared with those published by Zoetis (**Table 3**). When weaning weight ratio was the dependent variable to more accurately account for an animal’s rank in its contemporary group, WW GMX score was significantly predictive in the simple model with sex and random sire (0.08944±0.02057, p-value = 1.8e-05). When Milk GMX Score was compared to weaning weight ratio, the effect was not significant (p-value = 0.16). A model containing both GMX WW Score and GMX Milk Score provided a better fit to the data and estimated larger effects for the GMX Scores compared with the models using one GMX Score (**Table 4**). Calf phenotypes were plotted against dam’s GMX Score in **Figure 2**.

**Figure 2.**
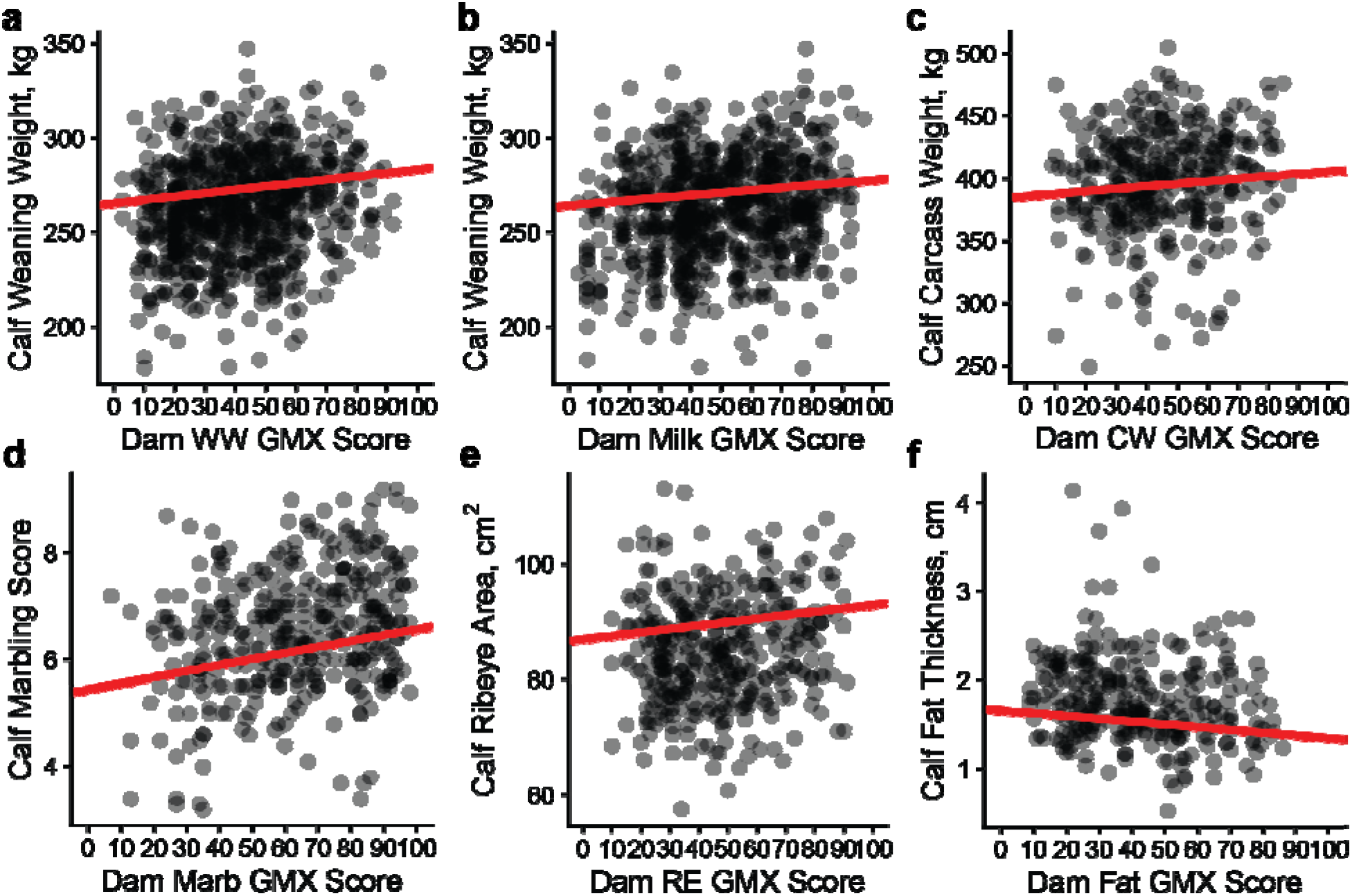
Calf’s phenotype plotted against dam’s GMX Score. a) Weaning weight versus WW GMX Score, b) Weaning weight versus Milk GMX Score, c) Carcass weight versus CW GMX Score, d) Marbling score versus Marb GMX Score, e) Ribeye area versus RE GMX Score, and f) Fat Thickness versus Fat GMX Score. Red line represents the intercept and slope estimated from the linear models reported in Table 3.

**Table 2.** Pearson and Spearman correlation tests between the dam’s Zoetis GeneMax Advantage Score (Kalamazoo, MI) and the calf’s phenotype (not adjusted for contemporary group or sire effects).

| GeneMax Trait | Pearson<br>Correlation<br>(95% Confidence<br>Interval) | Pearson<br>Correlation<br>P-value | Spearman<br>Correlation | Spearman<br>Correlation<br>P-value |
| --- | --- | --- | --- | --- |
| Weaning Weight | 0.18<br>(0.11 to 0.25) | 5.4e-07 | 0.18 | 3.0e-07 |
| Maternal Milk | 0.18<br>(0.11 to 0.25) | 2.5e-07 | 0.16 | 5.3e-06 |
| Marbling | 0.27<br>(0.18 to 0.36) | 8.4e-08 | 0.25 | 1.3e-06 |
| Carcass Weight | 0.13<br>(0.02 to 0.22) | 1.5e-02 | 0.15 | 4.8e-03 |
| Fat Thickness | -0.18<br>(-0.29 to -0.07) | 1.9e-03 | -0.19 | 1.1e-03 |
| Ribeye Area | 0.12<br>(0.02 to 0.22) | 2.0e-02 | 0.11 | 3.0e-02 |

**Table 3.** Estimated effects of GMX Scores on production traits. Each row represents a different linear mixed model. Models contained contemporary group and sex as fixed effects and sire as a random effect. Difference from zero *P*-values and MGS *P*-values are from *χ*^2^ test comparing full and reduced model. Difference from Zoetis effects estimated from a Z-test.

| GMX<br>Score | Estimate | Std. Error | GMX<br>effect<br>difference<br>from zero<br>*P*-value | Zoetis<br>effect | Difference<br>from<br>Zoetis *P*-<br>value | MGS<br>effect *P*-<br>value |
| --- | --- | --- | --- | --- | --- | --- |
| WW | 0.18 kg | 0.04 kg | 1.1e-05 | 0.25 kg | 0.08 | 1.1e-03 |
| Milk | 0.13 kg | 0.04 kg | 3.2e-04 | 0.14 kg | 0.88 | 1.1e-03 |
| Marb | 0.01 | 0.002 | 1.1e-06 | 0.01 | 0.21 | 3.2e-03 |
| CW | 0.20 kg | 0.09 kg | 3.3e-02 | 0.32 kg | 0.19 | 1.0* |
| RE | 0.06 cm<sup>2</sup> | 0.02 cm<sup>2</sup> | 2.2e-03 | 0.07 cm<sup>2</sup> | 0.58 | 9.5e-03 |
| Fat | -0.003 cm | 0.001 cm | 2.0e-02 | -0.03 cm | <1.0e-22 | 1.0* |

\*The estimated maternal grandsire variance for these models was zero.

**Table 4.**
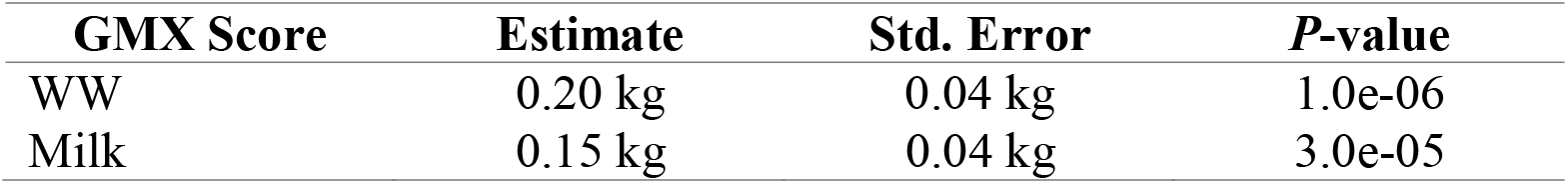
Estimated effects of dam’s WW and Milk GMX Scores on calf’s weaning weight. *P*-values from *χ*^2^ test comparing full model with both traits versus reduced models with single trait.

Tables 5 through 10 report additional model statistics. Models that included GeneMax score and sire fit the data better (lower AIC, lower BIC, higher pseudo-*R*^2^, better log likelihoods) than models that included maternal grandsire (MGS) and sire. Further, GeneMax scores were more significantly associated with progeny performance (smaller *P*-values) than the random effect of maternal grandsire.

**Table 5.**
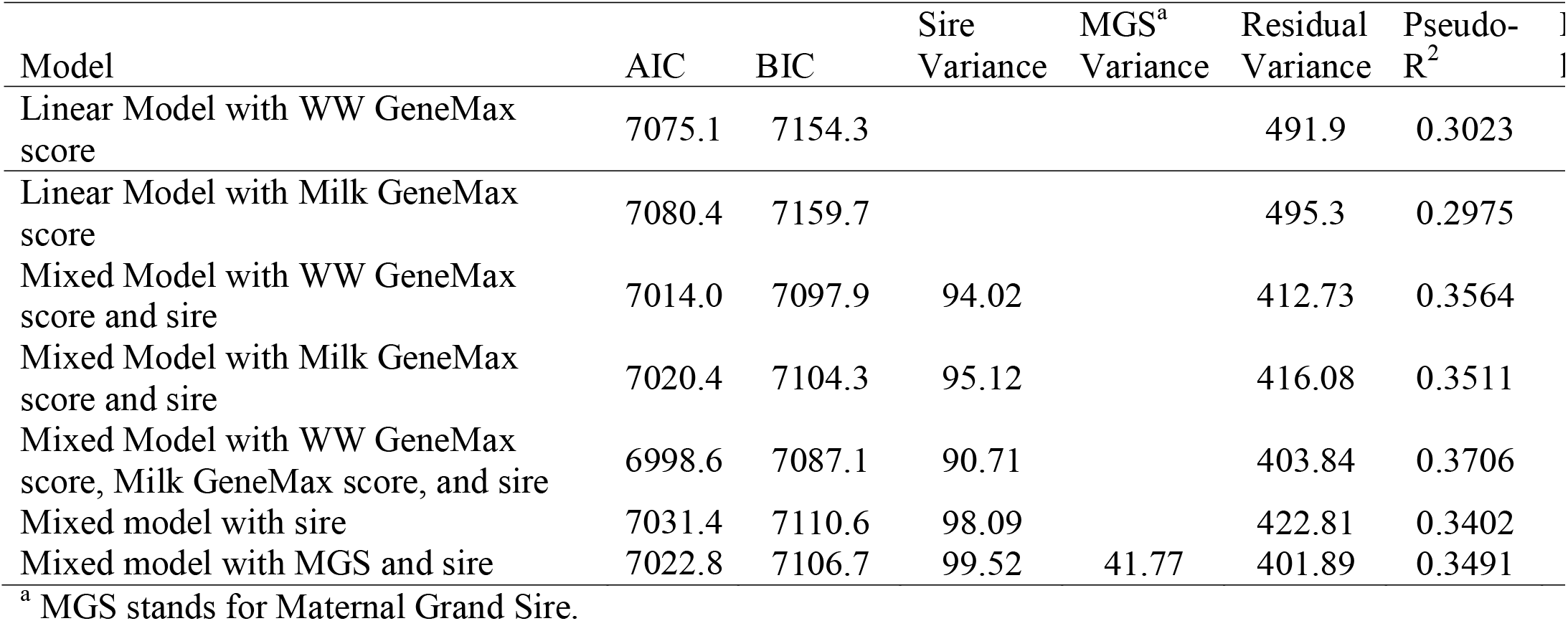
Summary of weaning weight models.

**Table 6.**
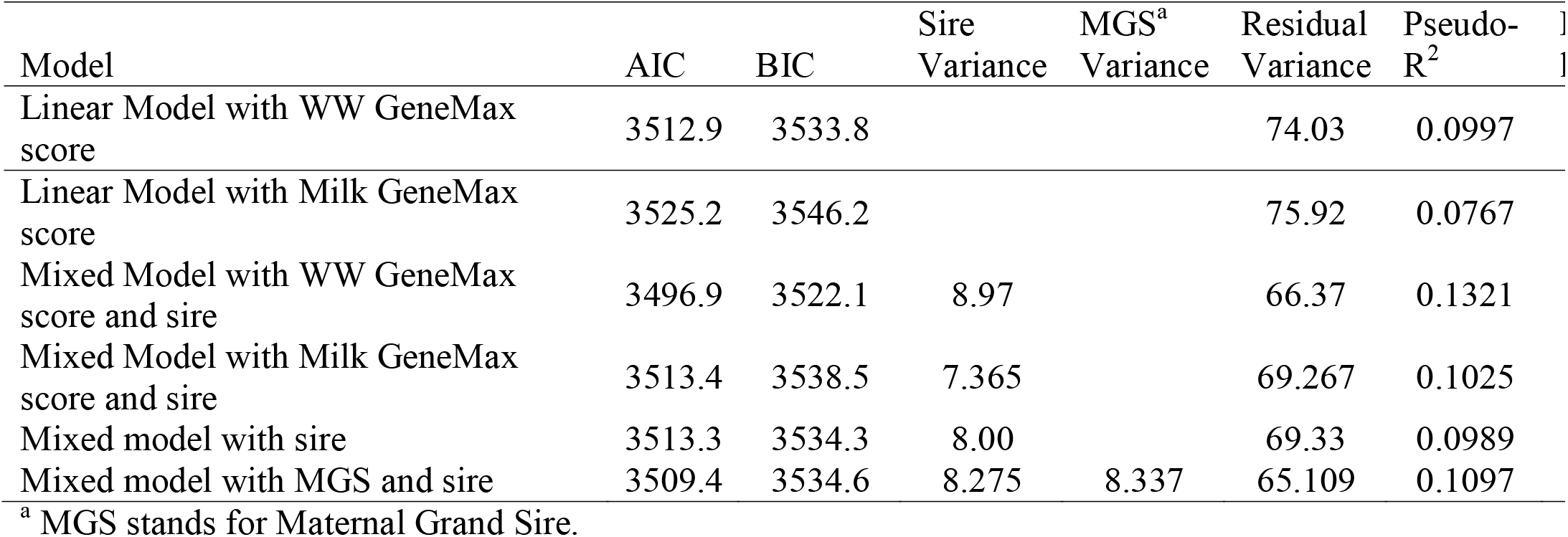
Summary of weaning weight ratio models.

**Table 7.**
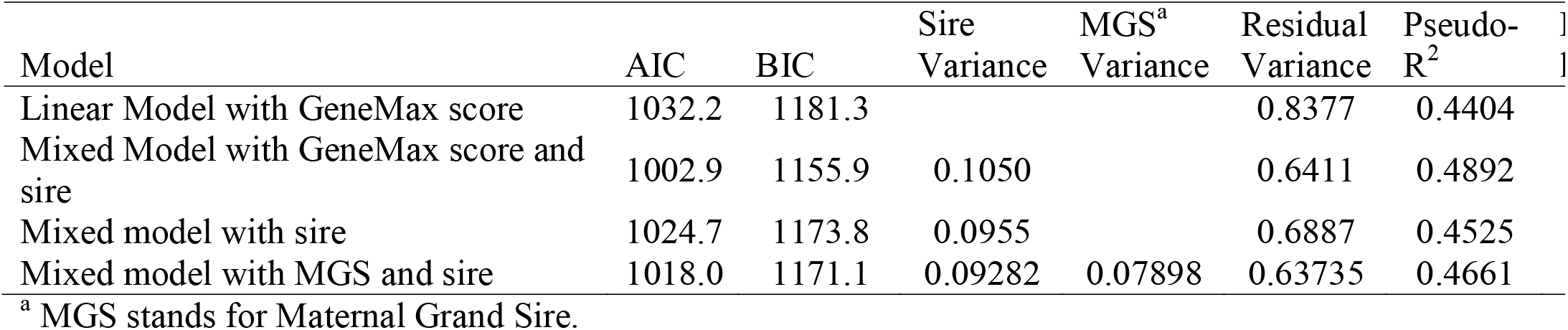
Summary of marbling models.

**Table 8.** Summary of carcass weight models.

| Model | AIC | BIC | Sire<br>Variance | MGS<sup>a</sup><br>Variance | Residual<br>Variance | Pseudo-<br>R<sup>2</sup> |
| --- | --- | --- | --- | --- | --- | --- |
| Linear Model with GeneMax score | 3724.2 | 3873.5 |  |  | 1062.5 | 0.4089 |
| Mixed Model with GeneMax score and sire | 3713.3 | 3866.6 | 161.9 |  | 848.3 | 0.4288 |
| Mixed model with sire | 3715.8 | 3865.2 | 152.1 |  | 862.0 | 0.4220 |
| Mixed model with MGS and sire | 3717.8 | 3871.1 | 152.1 | 0.0 | 862.0 | 0.4220 |

<sup>a</sup> MGS stands for Maternal Grand Sire.

**Table 9.** Summary of ribeye area models.

| Model | AIC | BIC | Sire<br>Variance | MGS<sup>a</sup><br>Variance | Residual<br>Variance | Pseudo-<br>R<sup>2</sup> |
| --- | --- | --- | --- | --- | --- | --- |
| Linear Model with GeneMax score | 2598.3 | 2747.4 |  |  | 55.17 | 0.4256 |
| Mixed Model with GeneMax score and sire | 2586.8 | 2739.9 | 7.105 |  | 44.329 | 0.4460 |
| Mixed model with sire | 2594.2 | 2743.4 | 6.418 |  | 45.747 | 0.4318 |
| Mixed model with MGS and sire | 2589.5 | 2742.6 | 7.952 | 5.821 | 41.880 | 0.4420 |

<sup>a</sup> MGS stands for Maternal Grand Sire.

**Table 10.** Summary of fat thickness models.

| Model | AIC | BIC | Sire<br>Variance | MGS<sup>a</sup><br>Variance | Residual<br>Variance | Pseudo-<br>R<sup>2</sup> |
| --- | --- | --- | --- | --- | --- | --- |
| Linear Model with GeneMax score | 369.65 | 432.04 |  |  | 0.1972 | 0.2527 |
| Mixed Model with GeneMax score and sire | 371.0 | 437.1 | 0.006747 |  | 0.1808 | 0.2552 |
| Mixed model with sire | 374.4 | 436.8 | 0.007184 |  | 0.1840 | 0.2347 |
| Mixed model with MGS and sire | 376.4 | 442.5 | 0.007184 | 0.00 | 0.1840 | 0.2347 |

<sup>a</sup> MGS stands for Maternal Grand Sire.

## Discussion

In the last ten years, the use of DNA information to produce genomic predictions has changed substantially. For example, when first launched in 2010, the IGENITY MBVs (molecular breeding values) were only based on 384 DNA markers (Weber et al., 2012). However, even the initial genomic predictions (which were use as indicator traits in a multi-step genomic-enhanced EPD analysis) trained with a couple thousand animals accurately predicted genetic merit (Weber et al., 2012). In the last ten years, hundreds of thousands of beef cattle have been genotyped from multiple breeds, increasing the power of these datasets not just for genetic prediction, but also for basic research (Decker, 2015). Since 2015, breed associations have switched to single-step methods, in which pedigree and genomic data are combined in a single analysis (Lourenco et al., 2015). Pedigree information is not typically known for commercial cattle, so a DNA marker effects model is typically used to predict genetic merit for commercial cattle. However, the estimated breeding values produced by a genomic relationship model and a marker effects model are equivalent (Hayes et al., 2009). The marker effects used to calculate GMX scores in the Zoetis GeneMax Advantage test are based on the American Angus Association single-step BLUP analysis (Zoetis Genetics and Angus Genetics Inc., 2018).

All traits had relatively weak correlations between the calf’s phenotype and the dam’s GMX Score. However, this is to be expected as this analysis does not account for Mendelian sampling (random shuffle of genes between generations), contemporary group effects (management and environment effects), sire effects, or the heritability of the trait. Nevertheless, as all correlations were significantly different from zero, it does demonstrate the predictive ability of the GeneMax Advantage test.

Regression analysis allowed a more sophisticated evaluation of the relationship between a dam’s GMX Score and her calf’s phenotype. These models accounted for variation due to sex, year of birth or slaughter date, and sire effects. However, this model still did not account for Mendelian sampling or other non-additive genetic effects, including genotype-by-environment effects. The amount of variation due to Mendelian sampling is large and theoretically equal to half of the additive genetic variance. These sources of variation are likely why we still observe substantial spread around the regression lines in **Figure 2**. Genetic predictions are not designed to predict performance of individual animals, but rather the average performance of a large group of progeny out of a parent compared to the progeny average of a different parent or population average. Our results show that the GeneMax Advantage test accurately predicts the average progeny performance for weaning weight, milk, marbling, carcass weight, ribeye area and fat thickness. Further, GeneMax scores provided more information than simply knowing the sire of the dam.

Our estimates of the effects of GeneMax scores did not differ from those published by Zoetis with respect to Milk (P = 0.88), Marb (P = 0.21), CW (P = 0.19), or RE (P = 0.58). However, our estimate of the effect of GeneMax score did differ from that published by Zoetis with respect to Fat (P < 1.0e-22) and tended to differ with respect to WW (P = 0.08). However, for all traits, our estimated effect sizes were smaller than those reported by Zoetis. These smaller effect sizes could be for many reasons, including data from a single environment in northwest Missouri, the size of the dataset, or that this was an external validation.

Regardless of whether we expressed weaning growth as a weight or as a ratio within contemporary group, WW GMX Score significantly predicted variation in weaning growth. However, Milk GMX scores were not predictive of weaning weight ratio. This likely reflects the well-known difficulty of predicting maternal effects (Willham, 1980). For example, in the 2014 Angus genomic-enhanced EPD calibration based on 57,550 animals, the correlation between weaning weight and Milk EPD was 0.36. The average of the other traits was 0.66 (range of 0.45 to 0.78). In 2016, when 108,211 animals were used to estimate molecular breeding values, the correlation between weaning weight and Milk EPD was 0.37, range of 0.56 to 0.80 for other traits (Albers, 2016). Further, only 491 observations were available when analyzing weaning weight ratio, compared to 781 observations for the models that only fit contemporary group, sex, and sire. Thus, the difficulty of predicting maternal effects and the smaller sample sizes affected the more complicated Milk GMX models.

We note that the Zoetis GeneMax Advantage prediction is designed to work in high-percentage Angus animals and is not designed for cattle with substantial ancestry from other breeds. However, other similar genomic predictions for crossbred cattle should be equally accurate provided they contain the appropriate breeds in a large, multi-breed training population (Kachman et al., 2013).

Genetic predictions, whether based on pedigree or genomic relationships, work when trained using ample and appropriately structured data. Models using contemporary group effects and random effects to account for covariance between relatives appropriately separate additive genetic variation from other sources of variation, including management and environment. While genetic predictions were never intended to predict the performance of a single individual, the average progeny performance is accurately predicted by the additive genetic merit of the parent. However, biological variation remains, including non-additive genetic effects and interactions (Smith et al., 2019; Braz et al., 2020) and especially Mendelian sampling (Cole and VanRaden, 2011) between full- or half-siblings. The increased adoption of genomic technologies in commercial cattle production has the opportunity to significantly increase long-term genetic gain through more accurate replacement animal selection.

## Conclusions

Genomic predictions, including the Zoetis GeneMax Advantage, accurately predict a straightbred, commercial Angus animal’s genetic merit and the average performance of their offspring. Academics and extension professionals can confidently state to farmers and ranchers that genomic predictions in commercial animals are accurate and effective.

## Abbreviations

GMX: GeneMax
SNP: single nucleotide polymorphisms
BIR: Beef Improvement Records
WW: Weaning Weight
MBV: molecular breeding value
BLUP: best linear unbiased prediction
CW: carcass weight
Marb: marbling
RE: ribeye area
Fat: fat thickness

## Acknowledgements

JED was supported by Agriculture and Food Research Initiative Competitive Grant numbers, USDA-NIFA 2016-68004-24827, USDA-NIFA 2018-68008-27891, 2020-67015-31132, and 2021-67021-33448 from the USDA National Institute of Food and Agriculture and USDA-NIFA Hatch MO-HAAS0027, USDA-NIFA MO-MSAS0014 (National Animal Genome Research Program). DJP was supported by USDA-NRI 2000-35203-9175, USDA-NRI 2005-55203-15750, USDA-NIFA 2007-55618-18238, USDA-NIFA 2013-68004-20364, and USDA-NIFA 2018-68008-27891. We acknowledge the support of the University of Missouri College of Agriculture, Food, and Natural Resources Thompson Research Center for providing labor, funds, and time for the collection of samples and data. This manuscript is published in memory of Miroslav Kaps. Miroslav Kapš was awarded his Ph.D. under the direction of William Lamberson at the University of Missouri in 1997 based on analysis of field records of Angus cattle. He was lead author of *Biostatistics for Animal Science*, a widely used reference text that was published in multiple editions and languages.

## Author Statement

**Brian. C. Arisman:** Formal analysis, Writing - Original Draft **Troy N. Rowan:** Formal analysis, Writing - Review & Editing, Visualization **Jordan M. Thomas:** Resources, Investigation **Harly J. Durbin:** Resources, Investigation, Supervision **William R. Lamberson:** Resources, Supervision **David J. Patterson:** Resources, Funding acquisition **Jared E. Decker:** Conceptualization, Methodology, Formal analysis, Writing - Review & Editing, Visualization, Supervision, Funding acquisition

## Literature Cited

Albers, C. 2016. Angus Announces Routine Calibration of GE-EPDs. Angus Media. Available from: http://www.angus.org/pub/newsroom/Releases/032116-recalibration.html

Barton K 2022. MuMIn: Multi-Model Inference. R package version 1.47.1, https://CRAN.R-project.org/package=MuMIn

Braz, C. U., T. N. Rowan, R. D. Schnabel, and J. E. Decker. 2020. Extensive genome-wide association analyses identify genotype-by-environment interactions of growth traits in Simmental cattle. bioRxiv. doi:10.1101/2020.01.09.900902. Available from: https://doi.org/10.1101/2020.01.09.900902

Cole, J. B., and P. M. VanRaden. 2011. Use of Haplotypes to Estimate Mendelian Sampling Effects and Selection Limits. J. Anim. Breed. Genet. 128:1–4. doi:10.1111/j.1439-0388.2011.00922.x. Available from: http://www.ncbi.nlm.nih.gov/pubmed/22059578

Decker, J. E. 2015. Agricultural Genomics: Commercial Applications Bring Increased Basic Research Power. G. Gibson, editor. PLoS Genet. 11:3. doi:10.1371/journal.pgen.1005621. Available from: http://dx.plos.org/10.1371/journal.pgen.1005621

Van Eenennaam, a L., J. Li, R. M. Thallman, R. L. Quaas, M. E. Dikeman, C. a Gill, D. E. Franke, and M. G. Thomas. 2007. Validation of commercial DNA tests for quantitative beef quality traits. J. Anim. Sci. 85:891–900. doi:10.2527/jas.2006-512. Available from: http://www.ncbi.nlm.nih.gov/pubmed/17178813

García-Ruiz, A., J. B. Cole, P. M. VanRaden, G. R. Wiggans, F. J. Ruiz-López, and C. P. Van Tassell. 2016. Changes in genetic selection differentials and generation intervals in US Holstein dairy cattle as a result of genomic selection. Proc. Natl. Acad. Sci. doi:10.1073/PNAS.1519061113.

Genetics, Z., and A. G. Inc. 2018. More Informed Commercial Angus Replacement Heifer Decisions with GeneMax^®^ Advantage^™^. Available from: https://www.zoetisus.com/animal-genetics/beef/pdf/zoetis-gmx-advantage-technical-bulletin-final.pdf

Hayes, B. J., P. M. Visscher, and M. E. Goddard. 2009. Increased accuracy of artificial selection by using the realized relationship matrix. Genet. Res. (Camb). doi:10.1017/S0016672308009981.

Kachman, S. D., M. L. Spanger, G. L. Bennett, K. J. Hanford, L. A. Kuehn, W. M. Snelling, R. M. Thallman, M. Saatchi, D. J. Garrick, R. D. Schnabel, J. F. Taylor, and E. J. Pollak. 2013. Comparison of molecular breeding values based on within- and across-breed training in beef cattle. Genet. Sel. Evol. 45:30. doi:10.1186/1297-9686-45-30. Available from: http://www.gsejournal.org/content/45/1/30

Lourenco, D. A. L., S. Tsuruta, B. O. Fragomeni, Y. Masuda, I. Aguilar, A. Legarra, J. K. Bertrand, T. S. Amen, L. Wang, D. W. Moser, and I. Misztal. 2015. Genetic evaluation using single-step genomic best linear unbiased predictor in American Angus. J. Anim. Sci. doi:10.2527/jas.2014-8836.

Meuwissen, T. H. E., B. J. Hayes, and M. E. Goddard. 2001. Prediction of total genetic value using genome-wide dense marker maps. Genetics. 157:1819–1829. Available from: http://www.pubmedcentral.nih.gov/articlerender.fcgi?artid=1461589&tool=pmcentrez&rendertype=abstract

Nejati-Javaremi, A., C. Smith, and J. P. Gibson. 1997. Effect of Total Allelic Relationship on Accuracy of Evaluation and Response to Selection. J. Anim. Sci. 75:1738–1745.

Smith, J. L., M. L. Wilson, S. M. Nilson, T. N. Rowan, D. L. Oldeschulte, R. D. Schnabel, J. E. Decker, and C. M. Seabury. 2019. Genome-wide association and genotype by environment interactions for growth traits in U.S. Gelbvieh cattle. BMC Genomics. 20:926. doi:10.1186/s12864-019-6231-y. Available from: https://bmcgenomics.biomedcentral.com/articles/10.1186/s12864-019-6231-y

Taylor, J. F., K. H. Taylor, and J. E. Decker. 2016. Holsteins are the genomic selection poster cows. Proc. Natl. Acad. Sci. U. S. A. 113. doi:10.1073/pnas.1608144113.

Team, R. D. C. 2018. R: A Language and Environment for Statistical Computing. R. D. C. Team, editor. R Found. Stat. Comput. 1. Available from: http://www.r-project.org

Weaber, R. L., J. E. Beever, H. C. Freetly, D. J. Garrick, S. L. Hansen, K. A. Johnson, M. S. Kerley, D. D. Loy, E. Marques, H. L. Neibergs, E. J. Pollak, R. D. Schnabel, C. M. Seabury, D. W. Shike, M. L. Spangler, and J. F. Taylor. 2014. Analysis of US Cow-Calf Producer Survey Data to Assess Knowledge, Awareness and Attitudes Related to Genetic Improvement of Feed Efficiency. In: 10th World Congress on Genetics Applied to Livestock Production. Vancouver, Canada.

Weber, K. L., D. J. Drake, J. F. Taylor, D. J. Garrick, L. A. Kuehn, R. M. Thallman, R. D. Schnabel, W. M. Snelling, E. J. Pollak, and a. L. Van Eenennaam. 2012. The accuracies of DNA-based estimates of genetic merit derived from Angus or multibreed beef cattle training populations. J. Anim. Sci. 90:4191–4202. doi:10.2527/jas.2011-5020.

Wickham, H. 2016. ggplot2: Elegant Graphics for Data Analysis. Springer-Verlag New York. Available from: http://ggplot2.org

Wickham, H. 2018. stringr: Simple, Consistent Wrappers for Common String Operations. Available from: https://cran.r-project.org/package=stringr

Wickham, H., R. François, L. Henry, and K. Müller. 2018a. dplyr: A Grammar of Data Manipulation. Available from: https://cran.r-project.org/package=dplyr

Wickham, H., and L. Henry. 2018. tidyr: Easily Tidy Data with “spread()” and “gather()” Functions. Available from: https://cran.r-project.org/package=tidyr

Wickham, H., J. Hester, and R. Francois. 2018b. readr: Read Rectangular Text Data. Available from: https://cran.r-project.org/package=readr

Willham, R. L. 1980. Problems in estimating maternal effects. Livest. Prod. Sci. 7:405–418. doi:10.1016/0301-6226(80)90080-9. Available from: https://linkinghub.elsevier.com/retrieve/pii/0301622680900809

Zoetis Genetics. (2016). Understanding and using GeneMax Advantage Results. Retrieved September 4, 2018, from https://www.angus.org/AGI/UnderstandingGMXAdvantageResults-08-26-16.pdf

